# Socru: Typing of genome level order and orientation in bacteria

**DOI:** 10.1101/543702

**Authors:** Andrew J. Page, Gemma C. Langridge

## Abstract

**Summary:** Genome rearrangements occur in bacteria between repeat sequences and impact growth and gene expression. Homologous recombination can occur between ribosomal operons, which are found in multiple copies in many bacteria. Inversion between indirect repeats and excision/translocation between direct repeats enable structural genome rearrangement. To identify what these rearrangements are by sequencing, reads of several thousand bases are required to span the ribosomal operons. With long read sequencing aiding the routine generation of complete bacterial assemblies, we have developed *socru*, a typing method for the order and orientation of genome fragments between ribosomal operons, defined against species-specific baselines. It allows for a single identifier to convey the order and orientation of genome level structure and 434 of the most common bacterial species are supported. Additionally, *socru* can be used to identify large scale misassemblies.

**Availability and implementation:** *Socru* is written in Python 3, runs on Linux and OSX systems and is available under the open source license GNU GPL 3 from https://github.com/quadram-institute-bioscience/socru.

## 1 Introduction

Bacterial genomes are dynamic entities that can undergo structural re-arrangement. These rearrangements tend to occur via homologous recombination around repeat sequences, including ribosomal rRNA operons and phage (Brüssow, et al., 2004; Sanderson and Liu, 1998). Different orders and orientations of large genome fragments have been described in many bacteria including *Enterobacter, Salmonella, Staphylococcus, Pseudomonas* and *Listeria* (Belda, et al., 2005; Chen, et al., 2017; Liu, et al., 2013; Tsuru, et al., 2006) while others appear to have a conserved genome structure e.g. *Klebsiella* (Ramos, et al., 2014).

To date, detection of structural rearrangements has been challenging, with low resolution methods such as restriction enzyme digestion and long-range PCR used to assay tens of strains at a time (Matthews, et al., 2011). The explosion of short read sequencing data over the past fifteen years has provided the necessary resolution for identifying small changes at the DNA level but consequently identifying structural variation at the whole genome level has lagged behind. However, the recent emergence of long read sequencing technology, which can bridge the length of long repeat sequences such as ribosomal operons, turns this situation around. As gross structural changes can impact upon growth and gene expression, knowledge of genome structure will improve the interpretation of results. We have developed *socru* for typing the order and orientation of genome fragments between ribosomal operons in complete bacterial assemblies

## 2 Implementation

A complete reference genome is required to provide a baseline order and orientation for a species. rRNA gene boundaries are identified using Barrnap (https://github.com/tseemann/barrnap), the nucleotide sequences (fragments) between the genes are extracted, circularized if they span the start/end and saved to individual FASTA files. By separating fragments into separate FASTA files, it allows for multiple representations of a fragment to be used, providing robustness in the method. Each fragment is compared to a database of *dnaA* nucleotide sequences using blastn, to identify the origin of replication, and this is noted in the database metadata.

The fragments are labelled numerically, beginning with the largest fragment and working in a clockwise fashion around the chromosome. Genome structures are represented using these numerical fragment numbers relative to the reference, with inverted orientations denoted with prime (‘). The genome structures always start with 1 and the orientation is always relative to the fragment with *dnaA* in the reference to provide consistency in the patterns.

To facilitate comparison of the overall structural variation in a population, each unique pattern is given a unique genome structure identifier (GS). The database contains a tab delimited table of these patterns. The genome structure identifier takes the form GSX.Y (for example GS4.16), where X uniquely denotes the order of the fragments and Y denotes the orientation of the fragments. The orientation is an integer representation of the orientation of the fragments in binary in reverse order (to allow for variability in the number of fragments), where 0 indicates same direction, and 1 indicates reverse direction. For example: 17’35642’ => 1000010 which is represented as GS2.<u>66</u>.

The software is bundled with a set of prepopulated databases covering 7401 genomes across 434 species. These represented the species with 3 or more complete reference assemblies available in RefSeq (accessed 2019-01-26), and where the reference sequence contained 3 or more rRNA genes. The assembly with the lowest numerical GCF accession number was chosen as the reference in each case. Patterns were accepted if they contained the same number of fragments as the reference, each occurring exactly once. The databases are stored on Github.com which allows for community curation and enhancements.

Given a FASTA file of a complete bacterial assembly, *socru* utilises a database (prebundled or user provided). First the location of the rRNA genes are identified with Barrnap. Each fragment is blasted against the user specified database of fragments, with a minimum e-value of 0.000001, minimum alignment length of 100 and minimum bit score of 100, and the top hit is used to identify the fragment number and the orientation. The order and orientation of the fragments is looked up in the bundled database of GS numbers. Novel orders are given a GS number of 0, which the researcher can evaluate for biologically probability. The output consists of the input file name, the GS identifier and genome structure pattern. The software requires less than 250MB of RAM to run and takes about 20 seconds to process a single 5Mbase assembly on a commodity laptop. The software is validated using unit tests and is packed for conda, galaxy, docker and pip for easy installation.

## 3 Application

We analysed all available complete *Salmonella enterica* genomes (n= 574) with *socru* using *S.* Typhimurium LT2 as the baseline for order and orientation. The genomes separated into 27 different GS types comprising 21 unique orders of which 6 had two differing orientations. The majority displayed the baseline order GS1.0 but around 20% displayed GS1.1 (inversion of fragment 1 relative to GS1.0) or GS2.67 (Fig. 1). For individual serovars of *Salmonella*, most displayed a single GS type but others showed greater variation: *S.* Enteritidis had 5 GS types and *S.* Newport had 3 types while at least one *S.* Typhi genome was represented in 17 GS types. Extended results of running *socru* on all complete genomes for the ESKAPE pathogens, *S. enterica* and *Escherichia coli* are available in Supplementary Table 1.

**Fig. 1.**
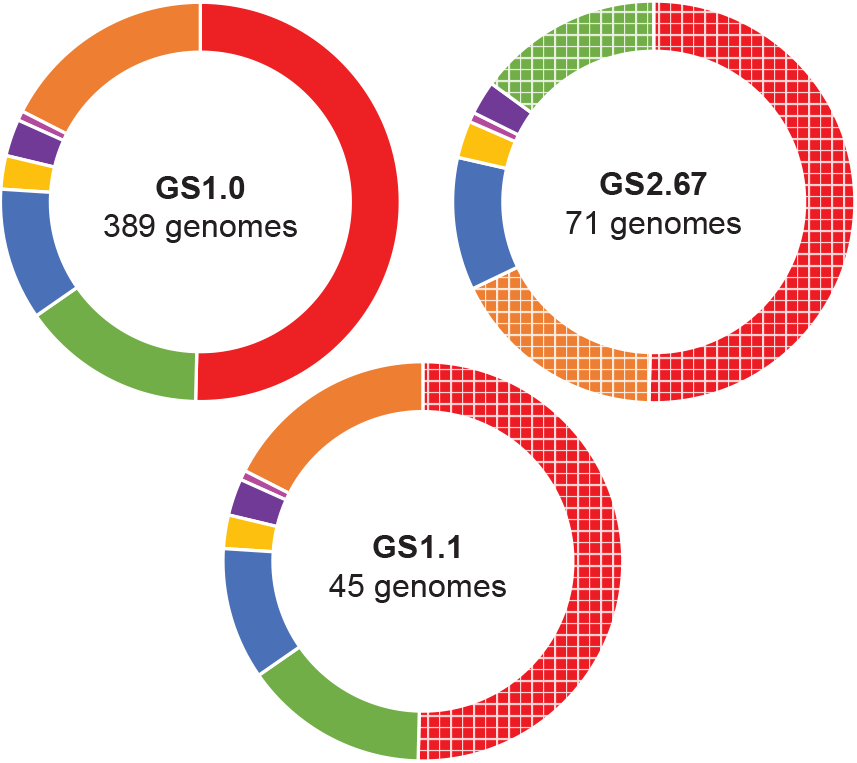
Common genome structures in *Salmonella*. GS1.0 represents the baseline order and orientation of the 7 genome fragments in *Salmonella enterica*. Inverted fragments (relative to baseline) in GS1.1 and GS 2.67 are indicated by hatched colours

## Acknowledgements

AJP was supported by the Quadram Institute Bioscience BBSRC funded Core Capability Grant (project number BB/CCG1860/1). GCL was supported by the Quadram Institute Bioscience BBSRC Strategic Programme: Microbes in the Food Chain (project number BB/R012504/1).

## Conflict of Interest

none declared.

